# Detection of HPV16, HPV18, p16, and E6/E7 MRNA in Nasopharyngeal Cancer: A Systematic Review and Meta-Analysis

**DOI:** 10.1101/401554

**Authors:** Rosalie Machado, Julian Khaymovich, Peter David Costantino, Tristan Tham

## Abstract

**Introduction:** Human Papilloma Virus (HPV) associated head and neck cancers, particularly oropharyngeal cancers (OPC), have a superior prognosis to HPV negative cancers. A literature review did not find any studies that analyzed HPV 16, HPV 18 DNA and p16 in the nasopharynx subsite, although it has been reported in the oropharynx. The detection of HPV DNA and p16 in the nasopharynx could have implications for the treatment of NPC. To date no one has reported the detection or concordance rates of HPV associated markers in nasopharyngeal carcinoma (NPC).

**Methods:** A literature search was undertaken on Medline, EMBASE, Scopus, and the Cochrane Library to identify studies that used PCR and ISH for detection of HPV DNA and p16 in the nasopharynx. We included studies published between 1992 and 2017 that reported the prognostic impact of HPV DNA and/or p16, treated HPV and p16 as categorical variables with at least 10 patients who tested for p16 and/or HPV16/HPV18 DNA in NPC, irrespective of HPV status. We then collected information from many parameters, and performed a meta-analysis to produce pooled prevalence estimates and explored sources of heterogeneity

**Results:** 21 studies published between 1992 and 2017 were selected for meta-analysis, with sample sizes ranging from 10 to 1328. The total (random effects) pooled detection of any HPV marker in NPC patients was 19.7% (95% CI: 12.56 to 28.00). The total (random effects) pooled detection of p16 positivity in NPC patients was 23.98% (95% CI: 14.82 to 34.54). The total (random effects) pooled detection of HPV DNA in NPC patients, detected by ISH, was 14.43% (95% CI: 10.13 to 19.33) and for HPV DNA in NPC patients including E6/E7 was 18.13% (95% CI: 11.45 to 25.94). The total (random effects) pooled detection of HPV DNA in NPC patients was 18.13% (95% CI: 11.45 to 25.94). The total (random effects) pooled detection of EBV positivity in NPC patients was 76.21% (95% CI: 66.40 to 84.80). Heterogeneity was high for all of the results. The attributable fraction of p16+ and HPV DNA+ in NPC is 0.029. The attributable fraction of EBV positive and HPV – in NPC is 0.827. The attributable fraction of EBV negative and HPV – in NPC is 0.0639. The attributable fraction of EBV negative and HPV + in NPC is 0.085. The attributable fraction of EBV positive and HPV + in NPC is 0.023. The attributable fraction of HPV positive and mRNA+ in NPC was 0.115. The attributable fraction of p16 in NPC was 0.123.

**Discussion:** This is the first study that contributes to the concordance and detection rates of HPV related markers in NPC.

**Limitations:** Most of the studies included were of Asian extraction. Most of the papers also did not test for E6/E7 RNA and therefore we are unsure of the amount of active E6/E7 RNA in NPC.

## Introduction

Nasopharyngeal carcinoma (NPC) is a malignant tumor arising from the epithelium of the nasopharynx often originating in the fossa of Rosenmuller and spreading outwards.^1^ Although, the cell type shares a common origin with other head and neck carcinomas, they have distinct differences. NPC is relatively infrequent; 86, 500 cases of nasopharyngeal carcinoma were reported worldwide in 2012, accounting for only 0·6% of all cancers.^1^ Further, the vast majority of the new cases (71%) were concentrated in eastern and southeastern Asia.^1^ Most of the rest were reported in south-central Asia, and north and east Africa.^1^ The incidence in Southeast Asia is 50/100,000 population compared to the United States which has an incidence rate in the single digits.^2^ This diversity is not limited to epidemiology, but extends to other clinical and biological features. The World Health Organization (WHO) has classified NPC into keratinizing (WHO type I) and non-keratinizing (WHO types II and III).^2^ The majority of the non-keratinizing carcinomas are in high-risk areas, while the keratinizing type is found mainly in low-incidence areas.^3^

The etiology of NPC is complex and involves the Epstein– Barr virus (EBV), genetic predisposition and other environmental risk factors such as smoking, salted-fish consumption and occupational exposures.^3^ Although, the exact oncogenic role of the EBV has not been determined, epidemiological evidence reveals that the preponderance of the undifferentiated NPC cases are EBV-related in intermediate-and high-incidence areas.^3^ Unlike the keratinized subtype, the differentiated and undifferentiated non-keratinizing carcinomas have raised Epstein– Barr virus titers.^3^ Several studies have demonstrated relationships between NPC and certain human leukocyte antigen (HLA) alleles and genetic polymorphisms.^3^

The presence of the EBV genome further subdivides NPC into groups with distinctly dissimilar clinical outcomes. The detection of nuclear EBV DNA in WHO types II and III, but not WHO type I NPCs, advocates that different pathogenic pathways operate for each tumor subtype.^2^ Kamran et al have linked the presence of EBV with NPC specifically in the endemic areas such as Southeast Asia.^4^

Fakhry et al investigated the role of Human Papilloma Virus (HPV) in the progression, local, regional, or distant, of oropharyngeal cancer (OPC).^5^ They found that it was limited in patients who had HPV positive OPC compared to those whose tumors were HPC negative.^5^ Studies have shown that HPV associated HNC, particularly oropharyngeal cancers, have superior prognosis to HPV negative cancers.^2^ This prognostic survival advantage has been pivotal enough to institute changes to the latest staging criteria of the AJCC into HPV+ and HPV-oropharyngeal cancer.^6^

Since the nasopharynx is anatomically contiguous with the oropharynx^7^ and they are both lymphoepithelial sites and share certain biological characteristics,^8^ the idea of a distinct disease entity, HPV-associated NPC, is an attractive one. HPV-associated or p16-associated NPC has been identified as a mutually exclusive infection from EBV infections by several authors.^9^

Overexpression of p16 in HPV oropharyngeal cancer indicates the inability of the body to control cell proliferation.^2^ In 2014, a meta-analysis by Ndiaye et al found that there was a significant concordance of HPV DNA and p16 in the oropharyngeal subsite.^10^ Over expression of p16 is associated with improved outcomes among patients with oropharyngeal cancer.^5^ However, researchers are unsure of a similar relationship with NPC. Recent studies have shown an oncogenic HPV associated with EBV negative NPC, whereas other studies have shown the HPV occurrence in NPC has no real prognostic value.^2^

The detection rates of HPV/p16 has been shown in the oropharynx and the oral cavity, but the literature has not reported the detection of HPV DNA and p16 in the nasopharnyx.^10^ A thorough review of the literature has shown that there have been no studies reported to date that analyzed HPV 16, HPV 18 DNA and p16 in the nasopharynx subsite, although it has been analyzed in the oropharyngeal and the oral cavity subsite.^10^ In addition, cancer databases such as National Cancer Database (NCDB) and the Surveillance, Epidemiology, and End Results (SEER) database does not differentiate between HPV DNA and p16 status and therefore it is not clear in these databases whether “HPV+” is defined by HPV DNA + or p16 +. This detection of HPV DNA and p16 has implications for the treatment of NPC. Since HPV+ OPC has been shown as having superior prognosis, the same prognostic effects might apply to NPC. Therefore, if indeed proven to be a prognostic factor, the detection rates of HPV DNA and p16 in NPC might influence the choice of treatment.

To the best of our knowledge, there have been no published reports about the global distribution and prevalence of HPV associated NPC. In this context, this systematic review and meta-analysis seeks to provide updated information about above, as well as to investigate the concordance rates of biomarkers of HPV-driven carcinogenesis such as p16INK4a and E6/E7 mRNA.

## Materials and Methods

### Design

Our search was performed in accordance with the Cochrane Handbook of DTA Chapter on searching [11]. We followed the Preferred Reporting Items for Systematic Reviews and Meta-Analyses (PRISMA) statement guidelines to identify, screen, and describe the protocols used in this systematic review. [12] [13] Our search strategy was designed in collaboration with a librarian at the Hofstra Northwell School of Medicine (WH), and the systematic review was prospectively registered in an online systematic review database (PROSPERO 2018:). [13]

### Search Strategy

Medline (via PubMed), EMBASE, Scopus and the Cochrane Library were searched on April 19, 2018. We searched all databases from their inception to the present and with restriction to only English as the language of publication and excluded any grey literature. To gather additional literature, bibliographies were hand searched and PubMed’s related articles search was performed on all included articles. Briefly, keywords used were variations of “cancer”, “carcinoma”, “malignancies”, “neoplasms”, “nasopharyngeal carcinoma”, “papillomaviridae”, “HPV”, “human papillomavirus” “p16”, and “Cyclin-Dependent Kinase Inhibitor p16.” In Cochrane, “Nasopharyngeal carcinoma” was excluded from search because the supplementary concept was not found in the Cochrane Mesh Search Builder. In PubMed, “cancer of nasopharynx” was removed from strategy because PubMed did not find the quoted phrase. In Embase, we filtered by document type to include articles, reviews, conference papers, short surveys, and articles in press. The full search strategy may be found in the supplementary materials.

### Article Selection

Articles were selected independently by two of the authors (TT, YB) in two phases. In the first phase we screened a list of titles and abstracts for full-text retrieval. During the first phase (title and abstract screening), our inclusion criteria was any study that reported a description of cancer of the nasopharynx, p16, and HPV, either in the title or abstract. If the content of the abstract was not clear, we selected the study for full-text review. Articles that passed the first phase of screening were selected for full-text retrieval, and were assessed in a second phase of screening.

In the second phase we screened full text articles using pre-determined inclusion/exclusion criteria.[13] Disagreements were resolved via consensus. For the second phase of the screening (full text retrieval), the following inclusion and exclusion criteria were applied. Inclusion criteria: (1) Article reports on prognostic impact of HPV DNA and/or p16; (2) HPV and p16 treated as categorical variables; (3) At least 10 patients tested for p16 and/or HPV16/HPV18 DNA in NPC, but not necessarily positive for HPV (4) Available as full text publication; (5) English language; (6) Clinical trial, cohort, or case-control study, letter to the editor. Exclusion criteria: (1) Case report, conference proceeding, reviews/meta-analyses; (2) Other types of head and neck tumors and Thyroid and endocrine tumors; (3) Animal studies; (4) Laboratory studies; (5) Duplicate literature and duplicate data, when multiple reports describing the same population were published, only the most recent or complete report to be included; (6) Incomplete data, although only excluded if contact with the original author does not provide missing information or if we are unable to reconstruct the data.

(1) Article reports on prognostic impact of peripheral blood NLR in head and neck cancer and associated subsites; (2) NLR treated as categorical variable; (3) NLR collected prior to treatment; (4) NLR Hazard Ratio (HR) / Risk Ratio (RR) for Overall Survival (OS), Disease specific survival (DSS), with or without Disease Free Survival (DFS), with or without Progression free survival (PFS); (5) 95% Confidence interval (CI) for survival statistic, with or without the p-value; (6) Available as full text publication; (7) English Language; (8) Clinical trial, cohort, case control. Exclusion criteria: (1) Case report, conference proceeding, letters, reviews/meta analyses; (2) Thyroid and endocrine tumors; (3) Animal studies; (4) Laboratory studies; (5) Duplicate literature and duplicate data; when multiple reports describing the same population were published, only the most recent or complete report was included; (6) Metastatic cancers only; (7) Incomplete data. Studies with incomplete data (for example, studies that included Kaplan-Meier curves only, or without HR with 95% CI), were not excluded initially.

The PRISMA flow chart of the systematic review can be found in Figure 1. An initial search done using the search strategy (Supplementary Materials) obtained an initial 1374 results. De-duplication was then performed, which reduced the number of results to 625. The first phase of screening was performed next on titles and abstracts, which reduced the number of results to 37. These 37 studies were additionally evaluated based on the inclusion/exclusion criteria, which resulted in a total of 21 studies. Thus, a total of 21 studies remained for quality assessment (supplementary materials).

**Figure 1:**
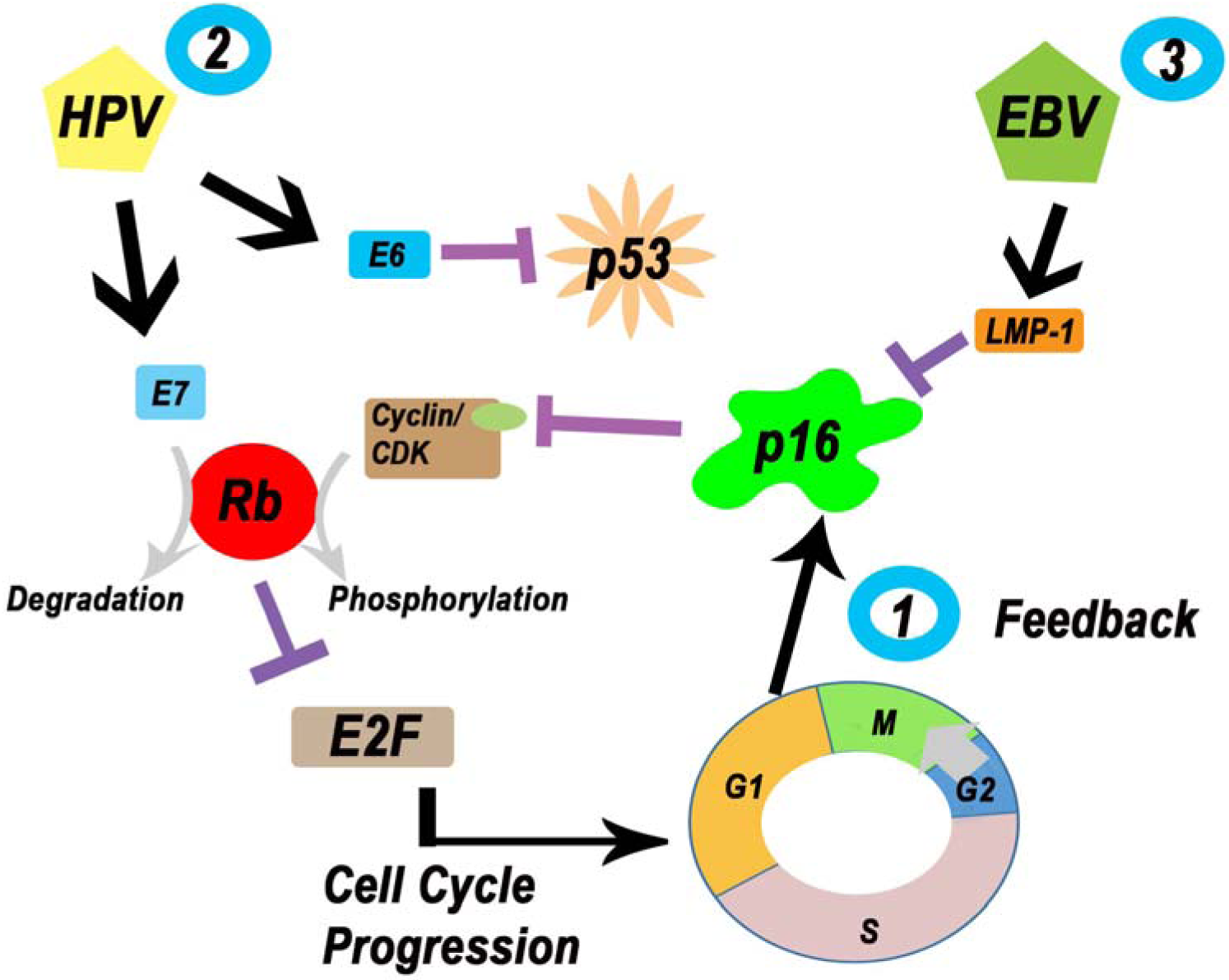

### Controlling for Methodological Heterogeneity

During quality assessment, we discovered there was significant methodological heterogeneity in the 37 initially selected papers. The papers were assessed for quality through the collection of eight criteria.^5^ These criteria were: (1) Mention of the use of PCR, (2) the use of human gene markers, (3) whether the cases were histologically confirmed, (4) mention of control for contamination, (5) inclusion of specific information on age and sex, (6) mention of recruitment period, (7) whether cases were recruited randomly or consecutively or were incident cases, and (8) whether sample size was at least 50 cases. The quality score was calculated as the sum of the results for each paper, where (0=no/1=yes) for all quality control criteria, making the lowest score zero and the highest score eight.

### Data Extraction

Data forms were developed *a priori* as recorded in the PROSPERO registry.[13] Two authors (TT, RM) jointly reviewed all of the full text articles together for the data extraction process. The following data points were collected: First author’s name; Year of publication; Country (region) of the population studied; Sample size; Age; Gender; type of study (prospective, retrospective, case-control, cohort, etc.); Study Population; Study Period; Journal of Publication; Case Selection Methods; Number of Nasopharyngeal cases available for pathological testing; Definition(s) of HPV positive status; demographic data; record of nasopharyngeal cancer; HPV detection, p16 detection, E6/E7 MRNA detection and EBV detection; Method of HPV DNA test; Method of p16 test; p16 threshold; Method of mRNA E6/E7; Method of EBV testing; Ethnicities; Smokers (ever/never); Alcohol consumption (ever/never); Tissue source (paraffin, fresh frozen); Total number of NPC cases tested for p16; and all HPV types tested E6/E7 mRNA+.

### Statistical Analysis

Pooled incidence was calculated for all the studies. Because of anticipated heterogeneity, a more conservative approach applying the random effects model (DerSimonian and Laird method) was chosen for all analyses. Forest plots were constructed for all outcomes displaying the random-effects model of the summary effect measure and 95% CI. Heterogeneity was assessed using Cochran’s Q and Higgins’s I^2^. Cochrane’s Q p-value of <0.1 and I^2^ > 50% were considered as markers of significant heterogeneity. All statistical tests were two-sided, and a p-value of less than 0.05 was considered statistically significant. MedCalc (Ostend, Belgium) was used for all statistical analyses.

## Results

### Study Characteristics

A total of 21 studies published between 1992 and 2017 were included in our meta-analysis, with sample sizes ranging from 10 to 1328 nasopharyngeal cases available for pathological testing. Our sample size was defined as the number of nasopharyngeal cases that were available for pathological testing. Seven studies were from the United States, three from Japan, two from China, and one each from Taiwan, India, Iran, Hong Kong, Mexico, and Belgium. Out of 21 studies, only one was a prospective cohort study. The rest of the studies were retrospective cohort studies. With regards to the outcomes reported, 15 of the studies reported any type of HPV marker including HPV 16 DNA, HPV 18 DNA, p16, or mRNA E6/E7, 14 of the studies reported p16 positivity, 20 studies reported HPV 16 or 18 DNA, and 14 studies reported EBV positivity.

### Detection of any HPV marker in Nasopharyngeal Cancer (NPC)

Data from 15 studies were synthesized in the meta-analysis for the detection of any HPV marker (HPV 16 and/or HPV 18 and/or p16 and/or mRNA E6/E7) in NPC patients. The total (random effects) pooled detection of any HPV marker in NPC patients was 19.7% (95% CI: 12.56 to 28.00, p<.00001). The test for heterogeneity showed an I^2^ value of 95.17%, p< 0.0001, which represented significant heterogeneity.

The detection of any HPV marker in NPC was stratified by endemic and non-endemic populations of NPC. For the non-endemic population, the total (random effects) pooled detection of any HPV marker in NPC patients was 20.46% (95% CI: 13.50 to 28.43, p<.00001). The test for heterogeneity showed an I^2^ value of 88.99%, p< 0.001, which represented significant heterogeneity. For the endemic population, the total (random effects) pooled detection of any HPV marker in NPC patients was 17.325% (95% CI: 3.42 to 38.81, p<.00001). The test for heterogeneity showed an I^2^ value of 93.81%, p< 0.001, which represented significant heterogeneity. The Forest Plot and corresponding Funnel Plot are represented in Figure 2.

### Detection of p16 in NPC

Data from 14 studies were synthesized in the meta-analysis for the detection of p16 positivity in NPC patients (Table 2). The total (random effects) pooled detection of p16 positivity in NPC patients was 23.98% (95% CI: 14.82 to 34.54, p<.00001). The test for heterogeneity showed an I^2^ value of 94.83%, p< 0.0001, which represented significant heterogeneity.

The detection of p16 positivity in NPC was stratified by endemic and non-endemic populations of NPC. For the non-endemic population, the total (random effects) pooled detection of p16 positivity in NPC patients was 23.83% (95% CI: 13.20 to 36.46, p<.00001). The test for heterogeneity showed an I^2^ value of 92.62%, p< 0.0001, which represented significant heterogeneity. For the endemic population, the total (random effects) pooled detection of p16 positivity in NPC patients was 24.79% (95% CI: 6.34 to 50.25, p<.00001). The test for heterogeneity showed an I^2^ value of 96.09%, p< 0.0001, which represented significant heterogeneity. The Forest Plot and corresponding Funnel Plot are represented in Figure 3.

### Detection of HPV DNA (HPV 16/ HPV 18) in NPC

Data from 20 studies were synthesized in the meta-analysis for the detection of HPV DNA positivity (HPV16/HPV18) in NPC patients (Table 2). The total (random effects) pooled detection of HPV DNA in NPC patients was 14.43% (95% CI: 10.13 to 19.33, p<.00001). The test for heterogeneity showed an I^2^ value of 83.94%, p< 0.0001, which represented significant heterogeneity. The total (random effects) pooled detection of HPV DNA in NPC patients was 18.13% (95% CI: 11.45 to 25.94, p<.00001). The test for heterogeneity showed an I^2^ value of 92.73%, p< 0.0001, which represented significant heterogeneity.

The detection of HPV DNA positivity in NPC was stratified by endemic and non-endemic populations of NPC. For the non-endemic population, the total (random effects) pooled detection of HPV DNA in NPC patients was 12.86% (95% CI: 8.13 to 18.49, p<.00001). The test for heterogeneity showed an I^2^ value of 78.71%, p< 0.0001, which represented significant heterogeneity. For the endemic population, the total (random effects) pooled detection of HPV DNA in NPC patients was 18.62% (95% CI: 7.88 to 32.61, p<.00001). The test for heterogeneity showed an I^2^ value of 89.55%, p< 0.0001, which represented significant heterogeneity.

The Forest Plot and corresponding Funnel Plot are represented in Figure 5.[67, 75-78]

Begg’s Funnel Plot for the detection of HPV DNA indicated that there was no evidence of publication bias. The *p* value for Egger’s test indicated that there was no publication bias for detection of HPV DNA in NPC patients (*p* =). A classic Failsafe *N* value was calculated which showed that additional——contradictory studies are needed to invalidate the current meta-analysis results of the detection of HPV DNA in NPC.

### Detection of EBV in NPC

Data from 14 studies were synthesized in the meta-analysis for the detection of EBV positivity in NPC patients (Table 2). The total (random effects) pooled detection of EBV positivity in NPC patients was 76.21% (95% CI: 66.40 to 84.80, p<.00001). The test for heterogeneity showed an I^2^ value of 94.07%, p< 0.0001, which represented significant heterogeneity.

The detection of EBV positivity in NPC was stratified by endemic and non-endemic populations of NPC. For the non-endemic population, the total (random effects) pooled detection of EBV positivity in NPC patients was 68.25% (95% CI: 57.76 to 77.88, p<.00001). The test for heterogeneity showed an I^2^ value of 86.34%, p< 0.0001, which represented significant heterogeneity. For the endemic population, the total (random effects) pooled detection of EBV positivity in NPC patients was 90.48% (95% CI: 82.62 to 96.15, p<.00001). The test for heterogeneity showed an I^2^ value of 84.71%, p< 0.0001, which represented significant heterogeneity.

The Forest Plot and corresponding Funnel Plot are represented in Figure 6.**[67, 75-78]**

## Discussion

To the best of our knowledge, there have been no published reports about the global distribution and prevalence of HPV associated NPC. In this context, this systematic review and meta-analysis seeks to provide updated information about above, as well as to investigate the concordance rates of biomarkers of HPV-driven carcinogenesis such as p16INK4a and E6/E7 mRNA.

We found 21 studies published between 1992 and 2017 that were selected for meta-analysis, with sample sizes ranging from 10 to 1328. The total (random effects) pooled detection of any HPV marker in NPC patients was 19.7% (95% CI: 12.56 to 28.00). The total (random effects) pooled detection of p16 positivity in NPC patients was 23.98% (95% CI: 14.82 to 34.54). The total (random effects) pooled detection of HPV DNA in NPC patients, detected by ISH, was 14.43% (95% CI: 10.13 to 19.33) and for HPV DNA in NPC patients including E6/E7 was 18.13% (95% CI: 11.45 to 25.94). The total (random effects) pooled detection of HPV DNA in NPC patients was 18.13% (95% CI: to 25.94). The total (random effects) pooled detection of EBV positivity in NPC patients was 76.21% (95% CI: 66.40 to 84.80). Heterogeneity was high for all of the results. The attributable fraction of p16+ and HPV DNA+ in NPC is 0.029. The attributable fraction of EBV positive and HPV – in NPC is 0.827. The attributable fraction of EBV negative and HPV – in NPC is 0.0639. The attributable fraction of EBV negative and HPV + in NPC is 0.085. The attributable fraction of EBV positive and HPV + in NPC is 0.023. The attributable fraction of HPV positive and mRNA+ in NPC was 0.115. The attributable fraction of p16 in NPC was 0.123.

We found a large heterogeneity within the methods used across the papers. Methods to detect HPV DNA on OPC cells are not reliable, complex and expensive and hence, immunostaining for p16 protein expression has been suggested as a substitute based on an established association between HPV positivity and p16 overexpression.^2^ p16, a cyclin dependent kinase inhibitor 2A, is a tumor suppressor protein that that thwarts unwanted cell proliferation. The HPV oncoprotein, E7, inactivates retinoblastoma (Rb) protein, an important cell cycle protein that serves to suppress the tumor and to control cellular proliferation.^2^ As a result of uncontrolled cell proliferation, p16 is released from negative feedback control and results in an increase in p16 levels in an attempt to control cell proliferation.^2^ (Figure 1)

In addition to delineating the HPV, p16, and EBV concordance rates for NPC, we suggest the following recommendation for future research. Future research must include detailed information on biomarker methods. Since there is a lack of information on transcriptional activity of HPV associated NPC, future studies might also want to investigate the concordance with E6/E7 mRNA. In addition to this, the prognostic effect of HPV in NPC is yet to be determined. Future studies should study both the HPV transcriptional activity as well as the relationship to survival outcomes in this group.

---

**Figure 1:**
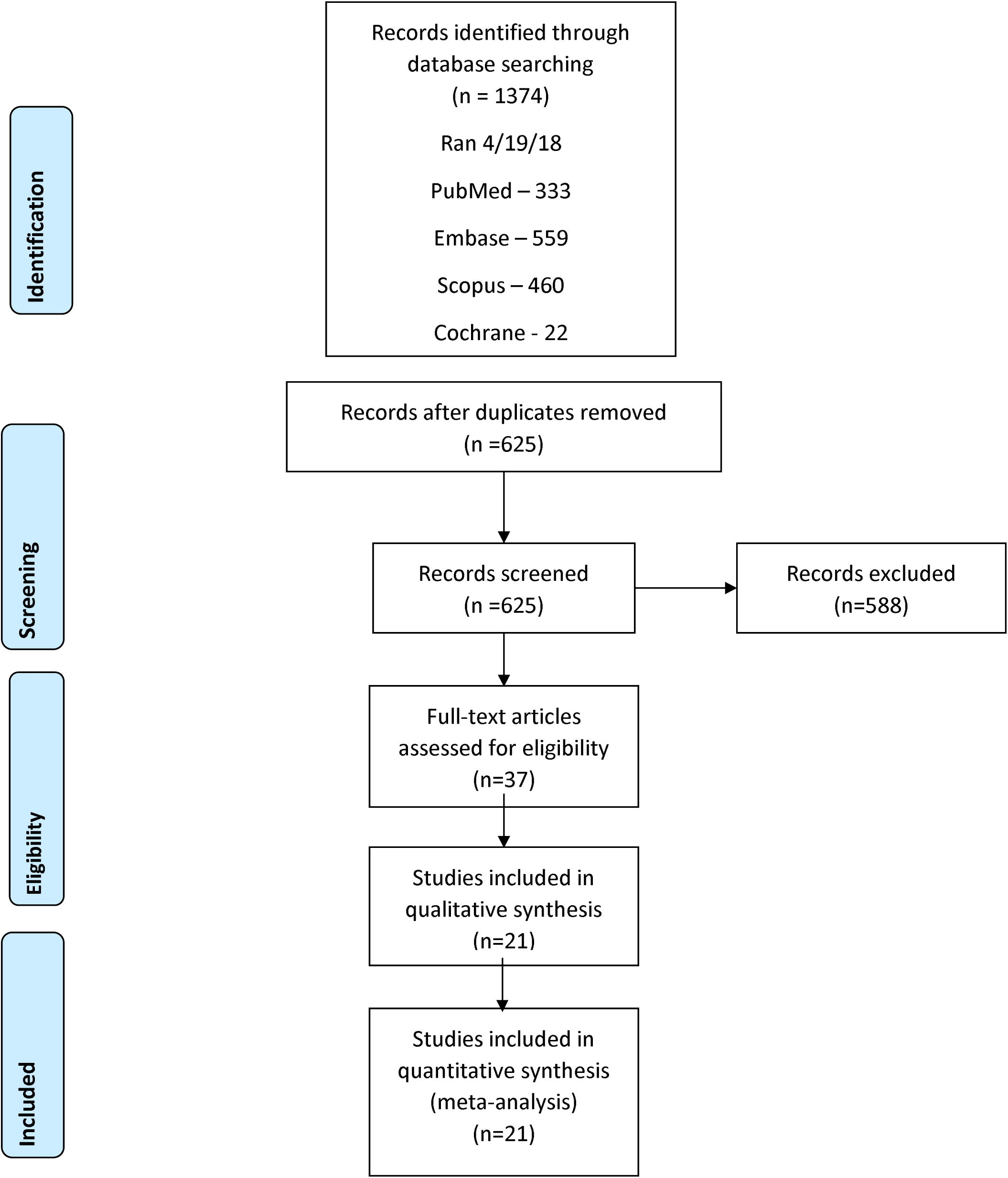
PRISMA.

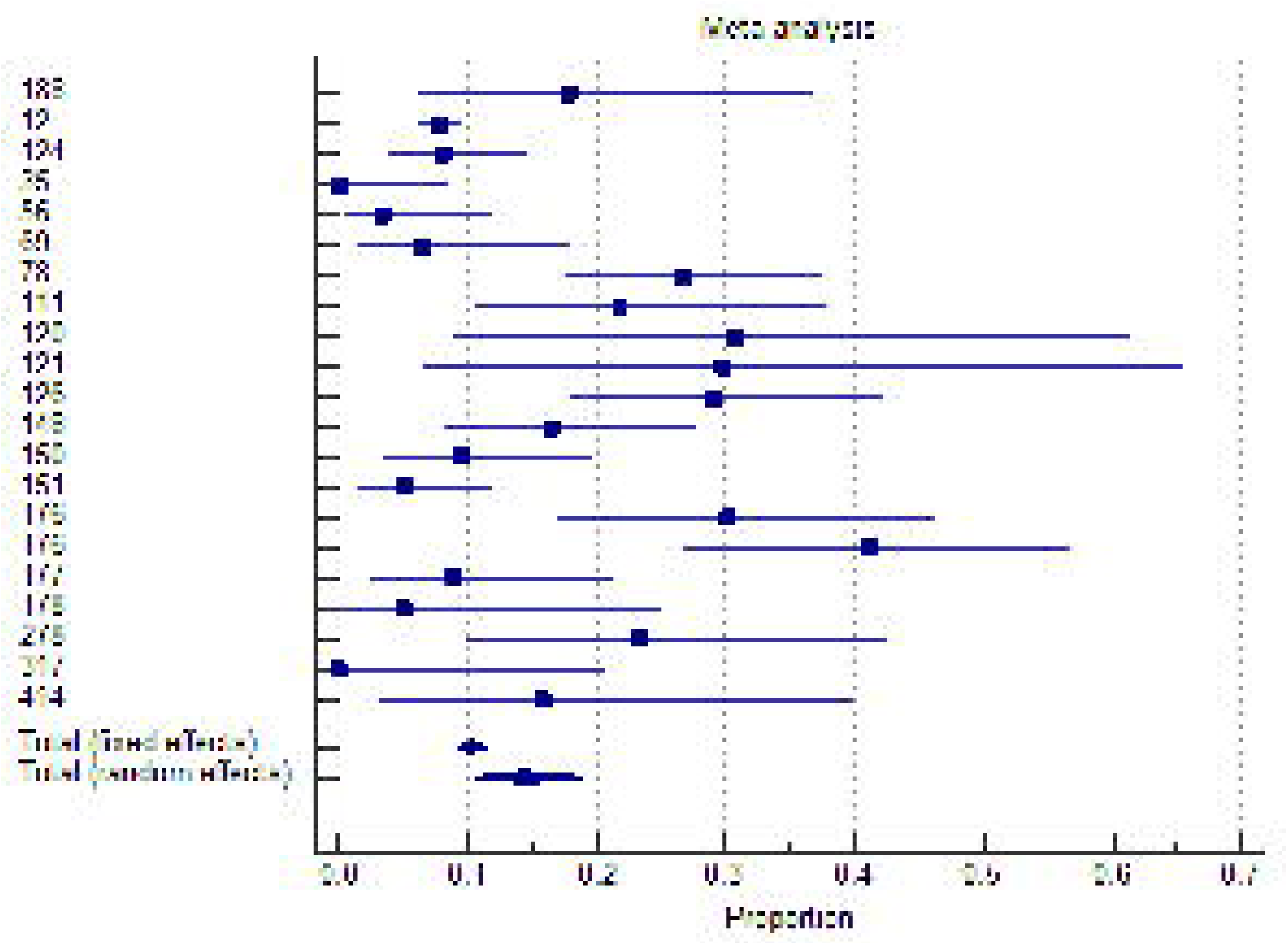

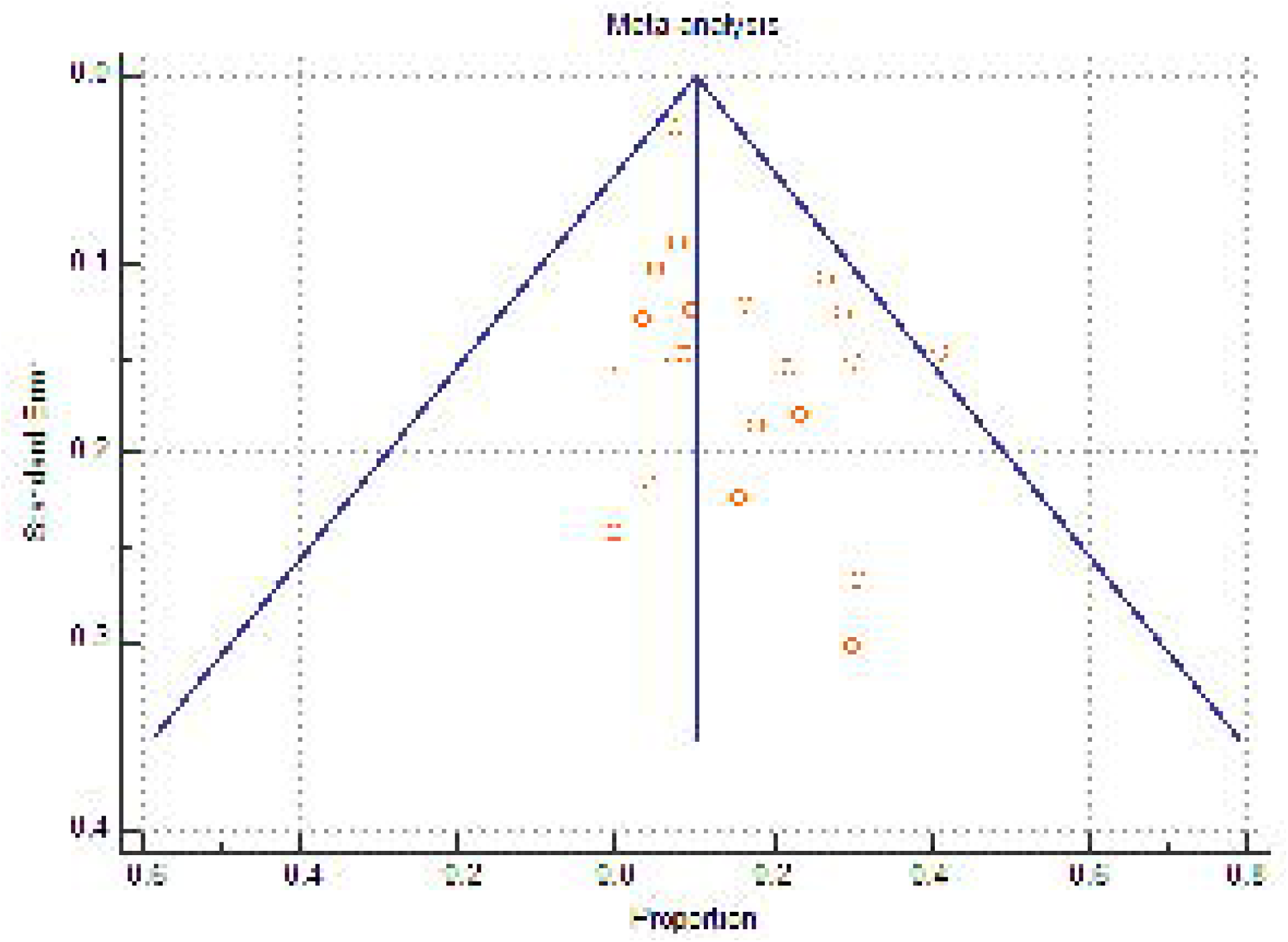

